# Modular metagenomic analysis of pan-domain symbioses with MAGUS

**DOI:** 10.64898/2025.12.22.696022

**Authors:** Gabriel A Al-Ghalith, Krista A Ryon, Erika Santoro, Adam Barno, Marco Casartelli, Helena Villela, Sara C Diana, James R Henriksen, Gaby E. Carpenter, Paola Quatrini, Marco Milazzo, Chirag J Patel, Raquel Peixoto, Braden T Tierney

## Abstract

Metagenomic analysis of deeply sequenced, eukaryotic-dominant symbiotic communities can be difficult for many metagenomic workflows. Here, we present MAGUS, a bioinformatic toolkit that uses a suite of custom bioinformatic methods for iterative genome assembly and filtering of pan-domain communities, where eukaryotes, bacteria, viruses, and functionally annotated gene catalogs are resolved and analyzed over a series of interconnected, modular software components. We evaluated MAGUS using deeply sequenced (median depth: 579 million reads) ten samples of hard corals, soft corals, and hydrozoans, which comprise complex, eukaryote-dominated symbiotic communities. We successfully resolved phylogenetically comparable host (N = 10), algal (N = 6), bacterial (N = 55), and viral (N = 160,925) genomes, as well as a gene catalog comprising 15,369,684 non-redundant genes (7.6% functionally annotated). MAGUS is available on GitHub (https://github.com/two-frontiers-project/2FP_MAGUS/).

## Main

Symbioses span the tree of life. In a given shotgun coral metagenome, 50-70% of sequencing reads derived from the host, 20-40% can map to its symbiotic algae, and the remainder will stem from bacteria, viruses, and other microbial eukaryotes.^1^ Lichens, legumes, leaf cutter ants, and numerous other holobionts house similarly non-monolithic communities.^2–4^ In recent years, the importance of assaying the complete diversity of metagenomic ecosystems has become apparent, with studies indicating the diversity of symbioses in nature and the role of organisms across the microbial tree of life in mitigating human disease and environmental dysbioses, ranging from heat-induced coral bleaching to pathogen colonization.^5,6^

“Ultra-deep” sequencing, where a single short-read sample contains hundreds of millions, if not billions, of reads, enables capturing genomic data from even low abundance organisms in a community, but it engenders numerous technical challenges^7^. High-quality genomes of rare taxa and microbial eukaryotes are underrepresented in public databases, so accurate reference-based alignment becomes challenging, and most bioinformatic tools are not optimized for eukaryotic genomes^8^. Reference-free approaches face additional difficulties: sequencing depth and within-sample diversity increase the computational resources needed to accurately assemble draft genomes, stretching assembly times to weeks or even months while risking creating chimeric contigs^9^. Co-assembly (i.e. combining replicate metagenomes into a single sample) can generate enough coverage to capture draft genomes from the low abundance symbionts, but as typically implemented it requires enormous computational resources and still carries the risk of chimeras due to abundant eukaryotic genomes^10,11^. As a result, there is a need for efficient tools that can resolve genomic diversity across the tree of life in deeply sequenced samples.

Here, we present a modular software suite, MAGUS, designed for the analysis of ultra-deep, shotgun sequencing of complex, multi-domain metagenomic communities via iterative assembly. It specifically is optimized for eukaryote-dominant metagenomes, but is relevant for any metagenomic analysis. In brief, MAGUS 1) subsamples reads, 2) takes a first-pass *de novo* assembly to identify putative, quality-controlled eukaryotic MAGs (based on a series of tools, including benchmarking on RefSeq eukaryotic reference genomes [see *Methods*])], 3) aligns reads against these MAGs to filter out the eukaryotic portion, 4) re-assembles the filtered dataset, 5) clusters these contigs to determine which samples should be co-assembled, and 6) runs a series of coassemblies. In particularly complex samples, filtering and binning eukaryotes can be repeated numerous times. From the resultant contigs at each stage of assembly, we provide numerous support functions (Supp Table 1) for binning bacterial, viral, and eukaryotic genomes, calling and annotating Open-Reading-Frames (ORF) and generating non-redundant gene catalogs, computing relative abundances with custom or public reference databases, genomic dereplication, and quality control.

MAGUS is not the first tool^12^ to tackle pan-domain metagenomic analysis, and it wraps numerous public tools, but it stands apart from previous efforts in its design and approach, most notably its ability to iteratively filter out and resolve bins from host reads. It also employs a novel and accelerated software tool for read alignment (XTree), assembly (megahit-g, which customizes megahit with new high-accuracy presets and adds the ability to handle longer k-mer sizes suitable for new sequencing technologies, like Ultima genomics systems, which is used in this manuscript)^13^, functional annotation (hmmsearch-g, which builds upon hmmsearch by overhauling the threading algorithm and disabling the bias filter)^14^, clustering and filtering (canolaX5, which wraps aKronyMer^15,16^ distances into an ultra-fast greedy clustering, read filtering, and pseudoalignment toolkit). Additionally, its approach to coassembly, identifying contigs not already binned into MAGs and shared between samples, enables an overall more efficient coassembly process geared towards novel MAGs. Finally, MAGUS is deliberately not a pipeline (i.e., a software workflow designed to be executed in linear or automated sequence), but rather an ecosystem of independent tools, where users are able to link analyses together in various orders to meet their needs (e.g., choosing to filter or not, building gene catalogs from MAGs or raw contigs, filtering bacteria or reads, etc). It is a flexible, modular pan-domain metagenomic analysis engine for any modern sequencing technology (e.g., short/long reads, single/paired-end reads).

We evaluated MAGUS’s functionality with ultra-deep, single-ended, sequencing of 10 different hard coral, soft coral, and hydrozoan fragments from the Red Sea that we collected on open-circuit SCUBA and sequenced for this study (see *Methods*). We specifically sampled species that did not have draft genomes at the time and were highly evolutionarily divergent. Metagenomic analysis of corals is a challenging task well-suited for MAGUS, as their genomes (average of 500Mb in size) are not well characterized, they have substantial genomic variation across species, and they contain symbionts with large genomes (up to 2,000Mb) from diverse clades. We sequenced samples with the Ultima Genomics UG100 sequencer to a mean depth of 579,690,672 (min/sample = 119,452,300, max/sample = 1,288,460,550), generating in total 5,796,906,725 reads (averaging 300bp each) of data. Our goal was to identify (1) draft coral and symbiotic algal genomes that could be phylogenetically characterized (i.e., meaningfully placed on a reference tree), (2) bacterial MAGs, (3) viral MAGs, and (4) non-redundant, annotated genes.

We processed each sample with the complete MAGUS workflow (see *Methods)*. This yielded species-level relative abundance data, contigs, bins (viral, eukaryotic, and bacterial), predicted open-reading-frames, and functional annotations for every stage of assembly. For the first pass assembly, samples were subsampled to 100 million reads. Both the filtered and the coassembly phases used all reads for a given sample that failed to align to the identified eukaryotic genomes in the initial stage.

As a first check, we measured the taxonomic composition of and total number of reads at each phase of assembly. Filtering yielded an average read count reduction of 83.5% (mean of 87,209,983 reads/sample) (Fig 2A). Contig N50 decreased, on average, from 3,134 base pairs to 1,610bp after filtering, due to the removal of large eukaryotic contigs, and total contig counts increased for hydrozoans and decreased for other species. We confirmed that bacterial reads were not being erroneously removed as the relative abundance of each species identified across all samples remained the same (pearson’s r = 1) before and after filtering.

**Figure 1:**
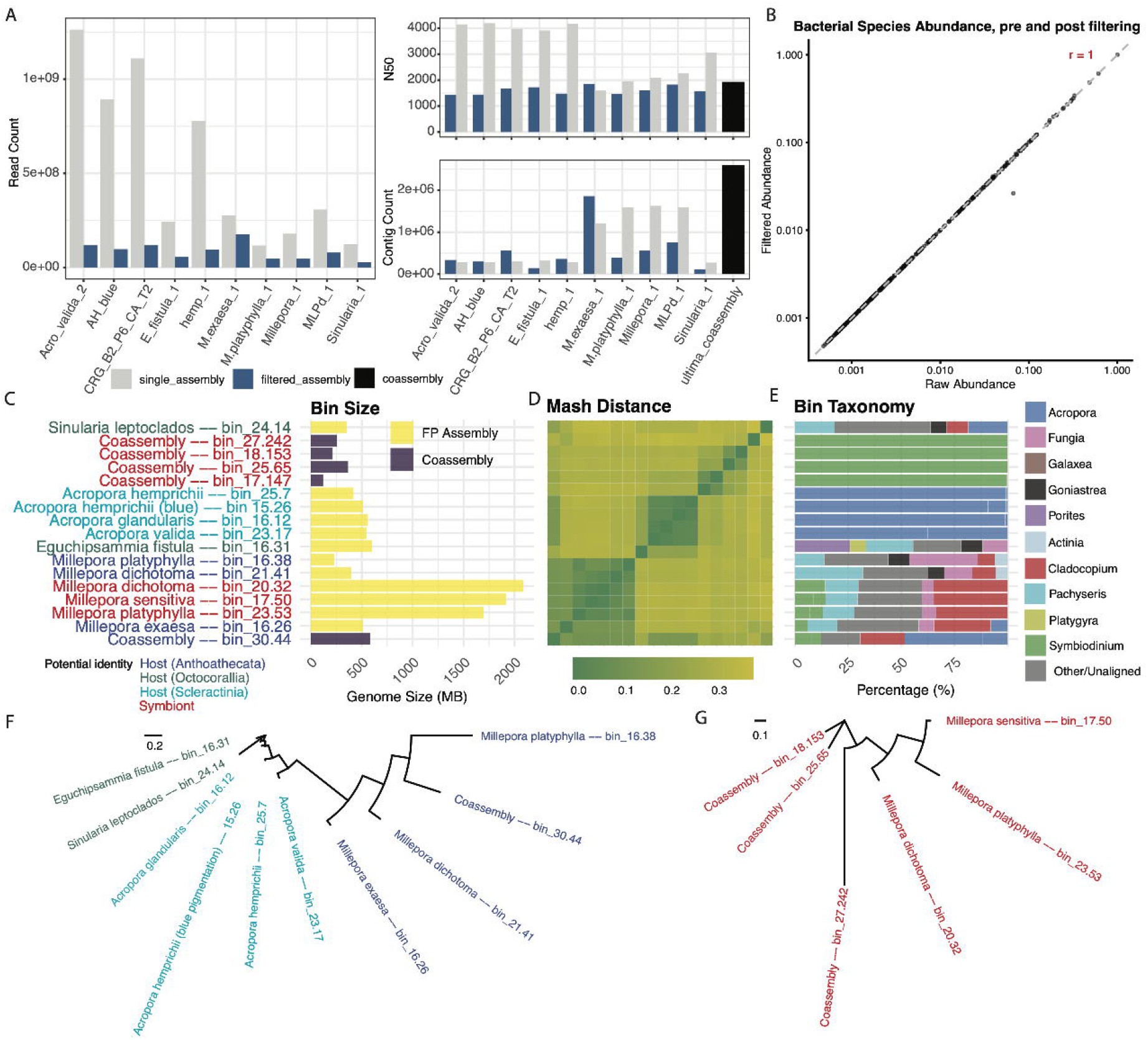
MAGUS filtering and identification of eukaryotic genomes in complex symbioses. A) Read count and contig summary statistics per sample analyzed for different phases of assembly (initial, filtered, coassembly). B) Abundance of bacterial species in samples before and after aligning to and filtering eukaryotic genomes. C) Sizes of putative eukaryotic bins D) Genome similarity between putative eukaryotic bins E) Taxonomy of contigs (based on alignment to Reef Genomics database) of all contigs in each putative eukaryotic bin. F) Phylogenetic tree of potential hydrozoan and soft/hard coral bins. G) Phylogenetic tree of potential algal symbiont bins.

**Figure 2:**
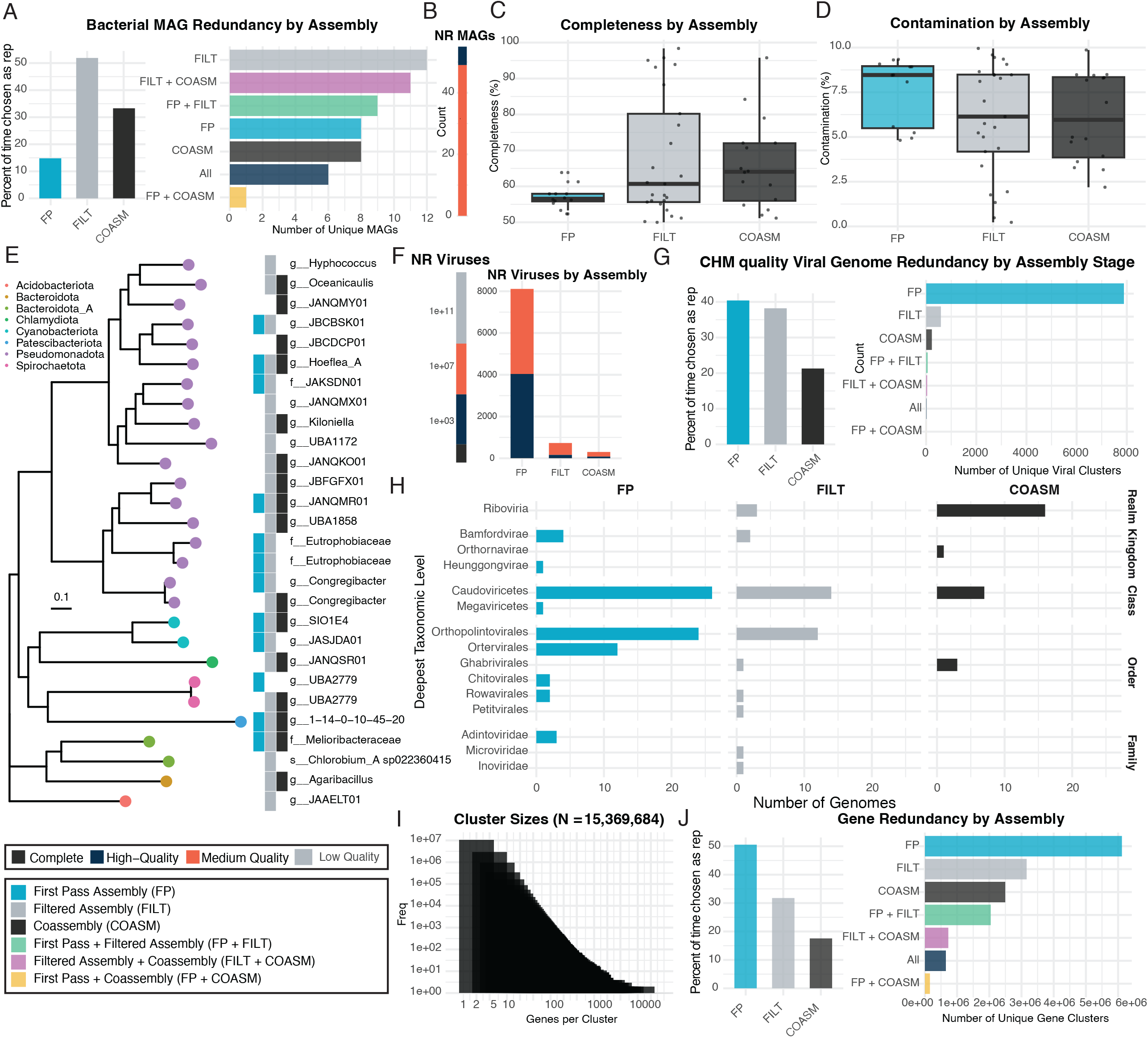
MAGUS captures bacterial, viral, and genetic diversity through iterative assembly. The color legend in the bottom left corner of the figure is to be used for all panels. A) Left panel: percentage of time a given bacterial MAG was identified as a representative per stage of assembly. Right panel: Number of unique bacterial MAGs identified by stage of assembly. B) Non-redundant (NR) bacterial MAGs by quality. C) Bacterial MAG Completeness by assembly stage. D) Bacterial MAG Quality by assembly stage. E) Phylogenetic tree of identified bacterial MAGs annotated by which assembly stage they were identified in. F) NR viruses identified, by assembly stage. G) Left panel: percentage of time a given viral genome was identified as a representative per stage of assembly. Right panel: Number of unique viral genomes identified by stage of assembly. CHM = Complete, High, Medium quality genomes. H) Viral taxonomies identified by assembly stage. I) Number of Open Reading Frames (ORFs) in each cluster in non-redundant gene catalog. J) Left panel: percentage of time a given ORF was identified as a representative per stage of assembly. Right panel: Number of unique genes identified by stage of assembly.

MAGUS resolved 17 non-redundant, putative eukaryotic genomes of sizes ranging from 120.8Mb to 2,089.2Mb (Fig 2C-E). Ideally, these would include at least one coral host genome for each sample and, potentially, a number of symbiotic algal genomes as well. 13 bins were found in the first pass assembly, and five came from the coassembly. Bin mash distances clustered by host phylogeny (i.e., hard corals clustered together, hydrozoan genomes clustered together, the one soft coral stood on its own). >99% of contigs from four bins (size from 420.0Mb to 561.0Mb) from *Acropora* samples aligned to *Acropora* references, indicating they were likely host genomes. *E. fistula* and *S. leptoclados* FP bins (601.1Mb and 353.5Mb, respectively) also aligned predominantly to corals, again indicating their likely identity as host genomes. One coassembled bin also appeared to be a partial host coral genome.

The *Millepora* samples, along with coassembly, resolved six putative algal symbiont genomes. 100% of the contigs in four of the coassembled bins aligned with equal confidence to *Sybiodinium* genomes, indicating their likely identity as algal symbionts. Three of the *Millepora* samples resulted in bins over 2,000Mb that clustered together based on mash distance and aligned to *Cladocopium* and *Symbiodinium*, indicating their likely identity as algal symbionts.

The other three, *Millepora-*derived bins (229.0Mb to 511.8Mb), we hypothesized were host genomes, but without further sequencing data (i.e., transcriptomics, long read sequencing), the nature of these genomes is difficult to ascertain due to the lack of existing *Millepora* references.

As an additional litmus test for our proposed taxonomic assignments, we called open reading frames (magus call-orfs –domain eukaryotic) and used ribosomal gene alignments to place putative hosts and symbionts on phylogenetic trees. The clustering of nodes aligned with the expected taxonomy of the genome in question. However, the smallest coassembly could not be placed on a tree due to lack of available markers. Despite this, though, we concluded that MAGUS enabled partial reconstruction of eukaryotic genomes of sufficient quality that they can, in general, undergo phylogenetic comparison.

For bacteria, MAGUS identified a total of 54, non-redundant medium to high quality (MIMAG standards) Metagenome-Assembled-Genomes (MAGS). The majority of representative species came from the filtered assemblies, followed by coassembly, followed by first pass assembly (Fig 3A), indicating both the benefit of increased sequencing depth for bacterial assembly without reduced risk of eukaryotic contaminants. We noted that the latter assembly stages identified a unique set of MAGs, indicating further benefits of accessing ultra-deep sequencing data and coassembly (Fig 3B). Completeness and contamination increased and decreased, respectively, after the first pass assembly (Fig 3C). For genomes of high-enough quality to be placed on a phylogenetic tree, only one had a species level annotation (Fig 3D). Identified taxonomies fell into *Psuedomonadota*, which is to be expected for corals.

Viral assembly, annotation, and dereplication yielded 151,779 non-redundant genomes (N complete = 21, N high-quality = 4,257, N medium quality = 4,868, N low quality = 151,779) (Fig 3F). 40.4% of representative genomes were selected from the initial assembly, with the remaining being identified in the later stages (Fig 3G, left plot). Most novel viruses were additionally selected during the first stage assembly pass (Fig 3G, right plot), however, coassembly accessed additional viral taxonomies beyond what was initially detected (Fig 3H).

Finally, MAGUS identified 15,369,684 non-redundant ORFs at the 30% amino acid identity level (Fig 3I). 7.6% of these were annotated with the Pfam-A HMM set. MAGUS can function with any HMM library and provides Kegg^17^ HMMs as well as Pfam-A^18^ as part of its release. The trends regarding novelty of genes followed the same pattern of viruses, with limited overlap between different assembly stages (Fig 3J).

Users can access MAGUS on GitHub (https://github.com/two-frontiers-project/2FP_MAGUS/). Dependencies are either accessible via conda or within the release repository’s bin itself. Future directions include expanding it to handling long read (co)assembly and additional functional annotation methods. Overall, we present this tool for use by the scientific community and look forward to its continued development.

## Figures and Tables

**Supplementary Table 1**: CheckM2 summary statistics on eukaryotic genomes from RefSeq, used to develop heuristics for identification of potential eukaryotic bins.

## Methods

### Overview of the MAGUS workflow

MAGUS employs iterative assembly to identify Metagenome-Assembled-Genomes (MAGs) from symbioses with members spanning the tree of life. First, to enable the first pass of *de novo* assembly, users can subsample all samples (magus subsample-reads) to a set, user-specified depth. This depth can potentially be determined with off the shelf, genomic diversity measurement tools like nonpareil^19^. Following subsampling, samples are assembled (magus assembly). Samples are then binned based on tetranucleotide frequency and bacterial MAGs are identified (magus binning). Eukaryotic bins can then be identified (magus find-euks). After aggregating all bins, they are dereplicated (magus dereplicate). We recommend aligning and filtering the individual from contigs back against the Genome Taxonomy Database to filter out potentially misbinned bacterial contigs. The identified eukaryotes are then indexed with XTree, reads are aligned against the resultant database (magus taxonomy), and filtered (magus filter-reads). Then binning can be run again, and filtering can continue iteratively until bin discovery tails off. Coassembly can additionally be run after any filtering stage (magus cluster-contigs followed by magus coassembly). Viruses can be identified also at any time (magus find-viruses). Gene catalogs are first constructed by calling Open Reading Frames (ORFs) with optional functional annotation by user-specific HMM databases (magus call-orfs). ORFs are then clustered into non-redundant gene catalogs (magus build-gene-catalog). For phylogenetic tree construction, we provide helper scripts to identify single copy genes and then build a tree with either FastTree or IQ-Tree^20,21^.

### Quality control

MAGUS qc wraps Shi7^22^ by calling shi7_trimmer on each sample with the preset arguments 75 12 FLOOR 4 ASS_QUALITY 20 CASTN 0 STRIP ADAP2 CTGTCTCTTATACA. It will function with either single or paired-end files. The trimmed FASTA outputs are immediately compressed with minigzip -4, and a tab-delimited post-QC configuration file recording the new FASTA locations is emitted for downstream steps.

### Read alignment and relative abundance calculation

The taxonomy module streams gzipped or plain reads into XTree ALIGN –redistribute --doforage. XTree output tables are merged and relative abundance is computed. Genomes are dropped based on a user-specific coverage value. When a taxonomy mapping (e.g., the Genome Taxonomy Database identifier to species mapping) is supplied, identifiers are harmonized and joined before pivoting to the final reference-by-sample matrix.

### Genome assembly

MAGUS assembly by default executes MEGAHIT (megahit-g) with a wide k-mer sweep (--k-min 75 --k-max 333 --k-step 6), stringent cleaning (--cleaning-rounds 1 --merge-level 100,.999), and a 1 kb minimum contig length for both single- and paired-end libraries. After assembly, contigs are renamed with a sample-specific prefix, and only those whose coverage metadata (multi tags ≥1.5 or ≥2) and length thresholds meet the heuristics are retained in temp0.fa for binning.

### Contig clustering and coassembly

MAGUS cluster-contigs first consolidates sample assemblies while excluding contigs drawn from large (>1 Gb) MAGs, trimming oversized files to maintain a 1 Gb per-sample cap. The remaining contigs are concatenated with lingenome, profiled with akmer102 (k=16, ANI/GC-local weighting), clustered by spamw2, and summarized with bestmag2 to generate a coassembly task list (coasm.todo) based on similarity between samples in terms of unbinned contigs. Coassembly groups concatenate raw reads within each cluster and re-run MEGAHIT using the same k-mer schedule, rename contigs to indicate the coassembly batch, and perform two rounds of coverage-informed filtering (≥1 kb and ≥500 bp) coupled with sorenson-g depth estimation and MetaBat2^23^ binning across seeds 15–30 (and offset seeds for the second pass).

### Eukaryotic binning

MAGUS find-euks searches user-provided bin directories, retaining only assemblies meeting the a user-specified size cutoff (10Mb is the default, and it was used in the analysis carried out in this manuscript). Each candidate is optionally screened with EukRep^24^ to identify eukaryotic contigs, then assessed with EukCC^25^ against the configured database using the requested thread count. CheckM2^26^ quality reports—either user-specified or auto-discovered—are normalized and merged so that completeness and contamination metrics can be appended to the final summary, which also records bin size, contig counts, and the number of EukRep-positive contigs.

For this manuscript, we also evaluated how CheckM2 classifies known eukaryotic genomes to develop heuristics for characterizing a bin as eukaryotic or otherwise (Supp Table 1). We downloaded all reference bacterial and eukaryotic genomes from RefSeq (N = 20,929 and N = 1,253, respectively) and ran CheckM2 with the default settings. We observed that CheckM2 contamination scores were most useful for characterizing something as eukaryotic, as it classified many eukaryotes as complete bacterial genomes (this could potentially be from high bacterial contamination in references, or the presence of symbionts). 99% of eukaryotes had contamination greater than or equal to 10% and completeness less than or equal to 100%. This captured 57% of all bacterial MAGs on RefSeq.

### Bacterial binning and taxonomic annotation

MAGUS binning computes coverage profiles by mapping reads to filtered contigs with sorenson-g -e 0.01, runs MetaBAT2 across a suite of seeds (15–30) to generate candidate bins, and evaluates them with CheckM2 (with optional custom databases). Bins are filtered using either explicit completeness/contamination thresholds or preset quality tiers (low/medium/high), copied into a “good” collection, and summarized via lingenome, akmer102 (k=13), spamw2, and bestmag2 to select representatives that are copied to the final MAG directory with provenance tracking. Taxonomic assignments for bacterial/archaeal, viral, and eukaryotic MAG FASTA collections are then produced by aligning each set with XTree against the appropriate database, capturing reference (.ref), coverage (.cov), and per-query (.perq) outputs and summarizing the top five species per contig using an auxiliary taxonomy mapping file.

### Viral binning

MAGUS find-viruses aggregates contigs from assemblies, appends unique identifiers to headers, and filters them by length bounds (default 500 bp–1 Gbp) before running CheckV^27^ end_to_end against the specified database. Quality summaries are filtered to the requested CheckV tiers (Complete/High/Medium/Low) while retaining the source identifier, and the resulting contigs are dereplicated with canolax5 (-k 14 -fitC 0.02). Representative status is propagated back into the filtered quality table to flag the final viral bin set.

### Sequence dereplication

MAGUS dereplicate symlinks all matching MAG files into a temporary directory, concatenates them with lingenome (HEADFIX mode), optionally trims any genome exceeding the user-specified maximum (default 2 Gbp), and clusters sequences with canolax5 using k-mer size 16, local alignment, and -fitC 0.05. Representative genome identifiers from the Canolax log are used to copy the full-length originals into an output directory while noting any genomes that were truncated for downstream awareness.

### Open-Reading-Frame calling, functional annotation, and gene catalog construction

MAGUS call-orfs differentiates by domain: bacterial MAGs run Prodigal in default mode, viral MAGs use prodigal-gv -p meta, eukaryotic genomes invoke metaeuk easy-predict with the user-selected database, and metagenomic contigs employ Prodigal’s metagenome model ^28–30^. For the eukaryotic ORF-calling in this study, we built a reference database from all coral and symbiont genes on UniProt. Output FASTA/FFN/GFF files are “manicured” to embed sample identifiers and sanitized characters, and optional hmmsearch passes (with configurable E-value cutoffs) create cleaned hit tables for downstream summaries. magus build-gene-catalog parses those summaries to split annotated from unannotated ORFs, clusters the latter with mmseqs2^31^ easy-cluster (--min-seq-id and coverage thresholds supplied by the user), and writes cluster, singleton, and single-copy tables that record sample IDs, gene IDs, and representative sequences.

### Identification of single-copy genes and tree construction

The phylogenomics helper scripts ingests a curated single-copy gene list and ORF summary, enforces a user-defined genome coverage threshold, and concatenates the best-scoring ORF per gene (falling back to placeholder sequences when absent). The single-copy gene list is generated by parsing the orf-calling output using another helper script provided within the MAGUS repository. The concatenated proteomes are aligned with MAFFT (--auto), optionally trimmed with trimAl using a user-specified gap threshold, and passed to either FastTree (default, multi-threaded via OMP_NUM_THREADS) or IQ-TREE (iqtree2) to infer the final phylogeny, while recording gene order information for reproducibility^20,21,32,33^.

### Collection of coral fragments

In December 2022, a subset of the coral samples was collected via SCUBA diving from the Coral Probiotic Village (CPV) reef complex in the central Red Sea (22°18′18.4″N, 38°57′52.5″E), following standard sampling ans storage procedures ^34,35^ a long-term coral reef study site established by researchers at King Abdullah University of Science and Technology (KAUST)^36^. This site, Al Fahal Reef, is a mid-shore reef extending several kilometers and located at depths of approximately 8–10 m, approximately 15 km off the coast of Saudi Arabia. Colonies were visually assessed for overall health, and 3–4 cm fragments were excised from each selected colony using sterile shears and immediately placed into 4 oz Whirl-Pak bags underwater. The following species were collected at CPV: *Acropora glandularis, Acropora hemprichii, Acropora valida, Millepora dichotoma, Millepora exaesa, and Sinularia sp*. On Rose Reef, a smaller reef in the central Red Sea (22°18′22.8″N, 38°53′07.2″E) at depths of approximately 18–30 m, Millepora platyphylla was collected using identical procedures. Additionally, a fragment of *Eguchipsammia fistula* was obtained from an aquarium mesocosm maintained at the Coastal and Marine Resources (CMR) Core Lab at KAUST. All samples were flash-frozen in liquid nitrogen immediately after collection and stored at −80 °C until subsequent processing.

### DNA Extraction and Sequencing

High-molecular-weight (HMW) genomic DNA was extracted from coral fragments using a modified protocol based on the QIAGEN MagAttract HMW DNA Kit (Cat. No. 67563; Qiagen, Germany). All extractions were conducted under sterile conditions. Up to 250 mg of frozen coral tissue was homogenized in a 2.0 mL microcentrifuge tube containing a metal grinding ball (MP Biomedicals, USA), 20 µL Proteinase K (Cat. No. 19134, Qiagen, Germany), 100 µL DNA/RNA Shield (Cat. No. R1100; Zymo Research, USA), and 220 µL Buffer ATL (Qiagen, Germany). Samples were incubated overnight at 56 °C with shaking at 500 rpm. Following lysis, 10 µL lysozyme (12.5 mg mL^-1^) (Cat. No. 89833; Thermo Fisher Scientific, USA) was added, and samples were incubated for 10 min at room temperature. Lysates were clarified by centrifugation, and 200 µL of the supernatant was transferred for DNA purification following the manufacturer’s recommended MagAttract HMW DNA Kit protocol. DNA was eluted in 52 µL of nuclease-free water and stored at 4 °C until library preparation.

DNA concentration and purity were assessed fluorometrically using a Qubit 4.0 Fluorometer (Thermo Fisher Scientific, USA) using the Qubit 1X dsDNA High Sensitivity chemistry (Cat. No. Q33230; Thermo Fisher Scientific, USA). A total of 300 ng of purified DNA was used for single-end (SE) sequencing with 300 bp reads on the Ultima Genomics UG 100 platform (Ultima Genomics, USA). Library preparation followed Ultima Genomics’ recommended protocol for HMW DNA input compatible with UG 100 chemistry.

## Data Availability

MAGUS is available at https://github.com/two-frontiers-project/2FP_MAGUS. Sequence data from this study can be accessed for non-commercial use at https://portal.geoseeq.com/projects/magus-paper-data.

## Acknowledgements

We thank Ultima Genomics for supporting the sequencing in this work. This project was funded by the Two Frontiers Project, a 501(c)(3) non-profit research organization.

## Author Contributions

G.A.G and B.T.T. conceived of and developed the tool. K.A.R and B.T.T. collected the samples, and K.A.R. extracted them for sequencing. All other authors read and edited the manuscript and/or tested, evaluated, and provided feedback on MAGUS as it was developed.

